# Multiple antibodies identify glypican-1 associated with exosomes from pancreatic cancer cells and serum from patients with pancreatic cancer

**DOI:** 10.1101/145706

**Authors:** Chengyan Dong, Li Huang, Sonia A. Melo, Paul Kurywchak, Qian Peng, Christoph Kahlert, Valerie LeBleu, Raghu Kalluri

## Abstract

Exosomes are man-sized vesicles shed by all cells, including cancer cells. Exosomes can serve as novel liquid biopsies for diagnosis of cancer with potential prognostic value. The exact mechanism/s associated with sorting or enrichment of cellular components into exosomes are still largely unknown. We reported Glypican-1 (GPC1) on the surface of cancer exosomes and provided evidence for the enrichment of GPC1 in exosomes from patients with pancreatic cancer^1^. Several different laboratories have validated this novel conceptual advance and reproduced the original experiments using multiple antibodies from different sources. These include anti-GPC1 antibodies from ThermoFisher (PA5-28055 and PA-5-24972)^1,2^, Sigma (SAB270028), Abnova (MAB8351, monoclonal antibodies clone E9E)^3^, EMD Millipore (MAB2600-monoclonal antibodies)^4^, SantaCruz^5^, and R&D Systems (BAF4519)^2^. This report complements such independent findings and report on the specific detection of Glypican-1 on the exosomes derived from the serum of pancreas cancer patients using multiple antibodies. Additionally, the specificity of the antibodies to GPC1 was determined by western blot and Protein Simple analyses of pancreatic cancer cells and their exosomes. Interestingly, our results highlight a specific enrichment of high molecular weight GPC1 on exosomes, potentially contributed by heparan sulfate and other glycosylation modifications.

## Introduction

We previously reported on the detection of the heparin sulfate proteoglycan Glypican-1 (GPC1) on serum-derived exosomes in patients with pancreatic cancer and breast cancer^1^. GPC1 expression is elevated in several cancer types and associated with poor prognosis^6-8^, including in pancreatic cancer cells, and mouse and human pancreatic cancer^8-14^. Informed by genetic studies, GPC1 is proposed to control fibroblast growth factor (FGF) signaling in mouse brain development^15^, and may be critical for pancreatic cancer cell proliferation and VEGF-A induced pancreatic tumor angiogenic response^9,10^. The proteoglycan GPC1, GPI-anchored to the cell surface, acts as a co-receptor for several heparin binding growth factors, thereby implicating it in promoting tumor progression^10-13,16^

In patients with pancreatic cancer, we described the enrichment of GPC1+ exosomes in the circulation when compared to healthy controls^1^. Exosomes are nano-sized extracellular vesicles produced by all cells and abundantly found in the circulation^17^. The renewed interest in their biology was fueled by their potential as liquid biopsies for various pathologies, including cancer. Although there are likely distinct subpopulations of exosomes with various biological properties, exosomes are generally carriers of the nucleic acids and proteins that reflect their cell of origin^17^. As such, exosomes from the serum of patients emerged as an attractive approach to possibly detect cancerous lesions and monitor and predict outcome of cancer patients. Several studies have reproduced our original findings related to enrichment of GPC1 in cancer exosomes, including in breast cancer exosomes, colorectal, esophageal and pancreas cancer^2-5,8,18-20^ (**Table 1**). Additionally, most of these reports show that such enrichment of GPC1 in exosomes can serve as biomarker. Either using the identical methodology reported in the Melo *et al.* publication^1^, or other improvised techniques, GPC1 was consistently identified as enriched in the exosomes of patients with different cancer types, including pancreatic cancer.

Although the mechanisms governing exosomes biogenesis remain to be elucidated, a potential enrichment and sorting of certain proteins, mRNAs and microRNAs into exosomes is being recognized^21,22^. It is interesting to note that exosomes from pancreatic cancer cells specifically enrich for the high molecular with GPC1, suggesting a specific, yet unknown mechanism for such transport to the surface of the exosomes.

## Results

Here we report on the use of three anti-Glypican 1 antibodies for the specific detection of GPC1^+^ exosomes in the circulation of pancreas cancer patients (n=10) compared to healthy donors (n=9) and benign pancreatic diseases (BPD, n=2) using flow cytometry as a readout. The ThermoFisher antibody used in our initial report^1^ (PA5-28055) is commercially available. We also utilized the antibody SAB2700282 from Sigma, and the monoclonal antibody MAB8351 (clone 9E9) from Abnova. Both ThermoFisher and Sigma described the immunogen as a recombinant fragment of human GPC1 corresponding to a region within amino acids 200 and 558. Abnova describes the immunogen as recombinant protein corresponding to full length human GPC1. We report on the highly reproducible and strong correlation between samples evaluated with each of these antibodies (**Figure 1A-B**). Notably, all 10 patients with PDAC showed elevated circulating GPC1+ exosomes bound-beads, in contrast with the 9 healthy donors and 2 BPD samples (**Figure 1B-C, Table 2**). We also proceeded with analyses using the ThermoFisher PA5-24972 anti-GPC1 antibodies that were generated to the synthetic peptides, however we noted these antibodies did not reproduce results compared to the antibodies used in this study (**Table 3**). Further, when exosomes were collected using polyethylene glycol (PEG) based isolation kits, the accuracy of detection was compromised.

**Figure 1.**
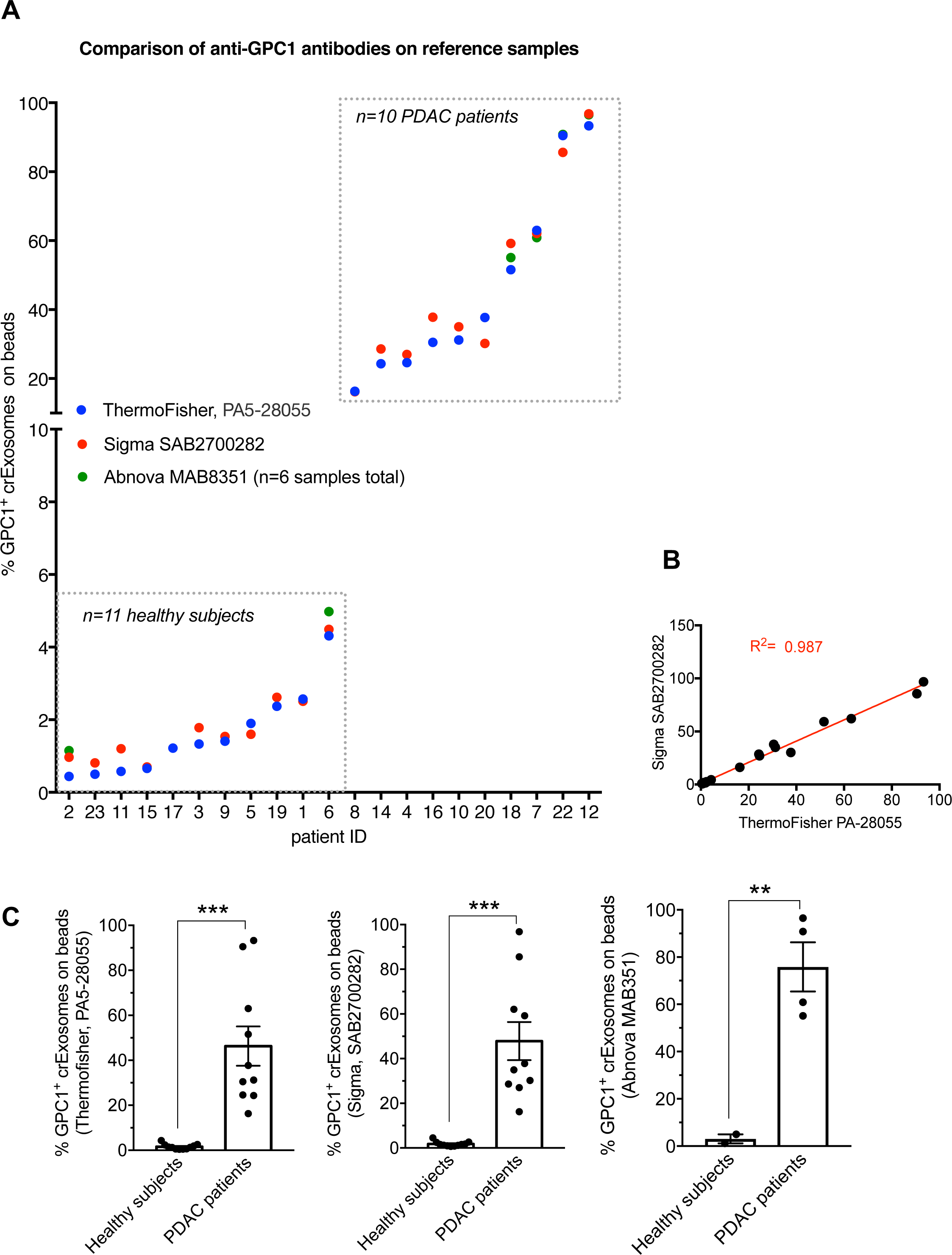
**A** Flow cytometry analyses for the % of GPC1+ circulating exosomes on beads, comparing the three anti-GPC1 antibodies listed. When the Abnova antibody was used, n=6 samples were analyzed (see Table 2). **B.** Linear regression analysis between % of GPC1+ circulating exosomes on beads obtained using the ThermoFisher PA5-28055 and Sigma SAB2700282 antibodies. **C.** Representation of the data shown in panel A, comparing healthy subjects vs. PDAC patients for three distinct antibodies (each dot represents a distinct patient/healthy subject). Unpaired two-tailed t test, ** p<0.01, *** p<0.001.

We carried out several experiments to inform on the specificity of the anti-GPC1 antibodies for the detection of cancer-derived exosomes. We utilized the human PDAC cell line Panc-1 (parental) and generated stable GPC1 knockdown (three distinct shRNA, referred to as shGPC1-1, shGPC1-2, shGPC1-3, see Methods) and over expression of GPC1 (GPC1 OE) clones. Sequence scrambled shRNA was used as control (shScrbl). All of the 3 different shGPC1 knockdown cell lines showed downregulation of GPC1 mRNA, while over expression of GPC1 was associated with upregulation of GPC1 mRNA, when compared to parental and shScrbl control cells (**Figure 2A**). Subsequently, several experiments were carried out on these cells and the exosomes derived from these cells.

**Figure 2.**
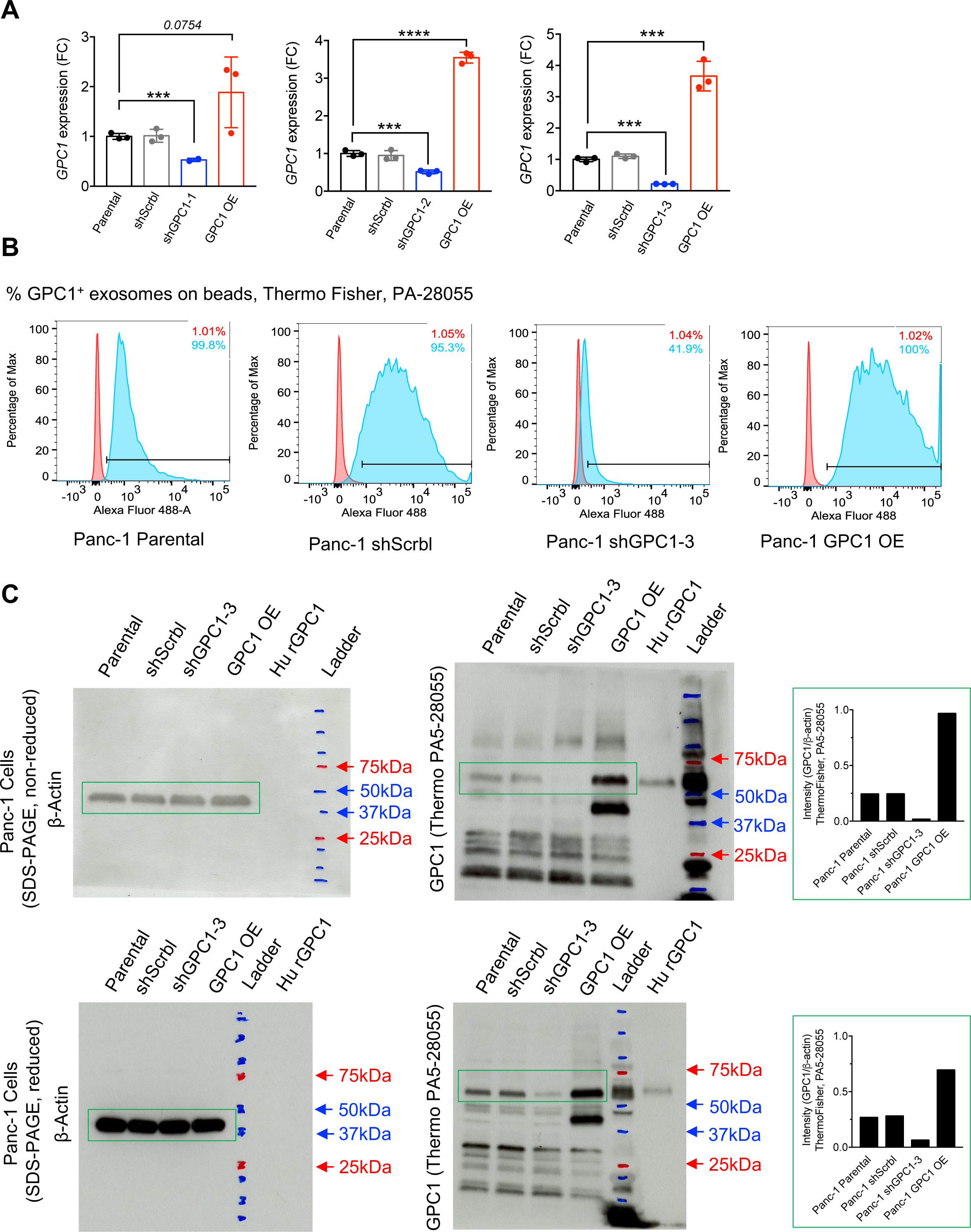
**A**. qRT-PCR measurement of GPC1 mRNA levels in Panc-1 cells (mean ± SEM; n=2-3). **B**. Flow Cytometry of the percentage of GPC1+ exosome on beads from Panc1 cells with GPC1 Primary Antibody (ThemoFisher PA5-280055). Negative control: red peaks, secondary antibody only. **C.** Western blot analyses of non-reduced and reduced Panc-1 cell lysates for GPC1 using ThemoFisher PA5-28055. β-actin is used as loading control. Recombinant human Glypican-1 protein (R&D Systems, 4519-GP-050) was used as positive control (Hu rGPC1). The graphs depict the quantification of GPC1 protein levels (densitometry analysis) relative to β-actin as captured in the green boxes.

As previously described and validated by others, exosomes were allowed to bind the beads and evaluated by flow cytometry for GPC1 positivity using the ThermoFisher PA5-28055 anti-GPC1 antibodies. Expected positivity for the *%* GPC1+ exosomes associated with the beads was noted in parental, shScrbl, and GPC1 OE cells-derived exosomes, and a decrease in the shGPC1 cells-derived exosomes (**Figure 2B**). Western blot analyses of non-reduced (as reported in Melo *et al.* publication^1^) and reduced cell lysates using the ThermoFisher PA5-28055 anti-GPC1 antibody revealed the specific loss of the core GPC1 protein (~60kDa), verified by the identification of the human recombinant GPC1 protein, in the shGPC1 cells compared to parental and shScrbl control cells (**Figure 2C**). In contrast, a specific accumulation of GPC1 (~60kDa) was noted in the GPC1 OE cell lysates compared to parental and shScrbl controls, in both non-reduced and reduced protein analyses (**Figure 2C**). The cell lysates show lower amounts of high molecular weight (HMW) heparan sulfate GPC1 protein, but low molecular weight (LMW) core GPC1 protein (~60kDa) was more abundant.

When Panc-1 exosomes lysates were evaluated using the ThermoFisher PA5-28055 anti-GPC1 antibody, the HMW GPC1 protein was abundantly detected in the exosomes (**Figure 3A**). When the same exosomes lysate preparation was analyzed under reducing conditions, the LMW core GPC1 protein (~60kDa) appeared (**Figure 3A**). Notably, a specific loss of both LMW and HMW GPC1 was observed in shGPC1-derived Panc-1 exosomes compared to parental and shScrbl controls (**Figure 3A**). The parental, shScrbl and GPC1-OE cells-derived exosomes were highly enriched for GPC1. These results reproduced when Panc-1 exosomes lysates under reducing conditions were analyzed using Protein Simple Western analysis system (www.proteinsimple.com) (**Figure 3B**). The Protein Simple Western analysis also reproduced the results when Sigma SAB270028 anti-GPC1 antibody was used (**Figure 4B**). Additionally, the Sigma SAB270028 anti-GPC1 antibodies used on conventional western blot analyses also reproduced the results for both cell and exosomes lysates, as observed with the ThermoFisher PA5-28055 anti-GPC1 antibodies (**Figure 4A-B**).

**Figure 3.**
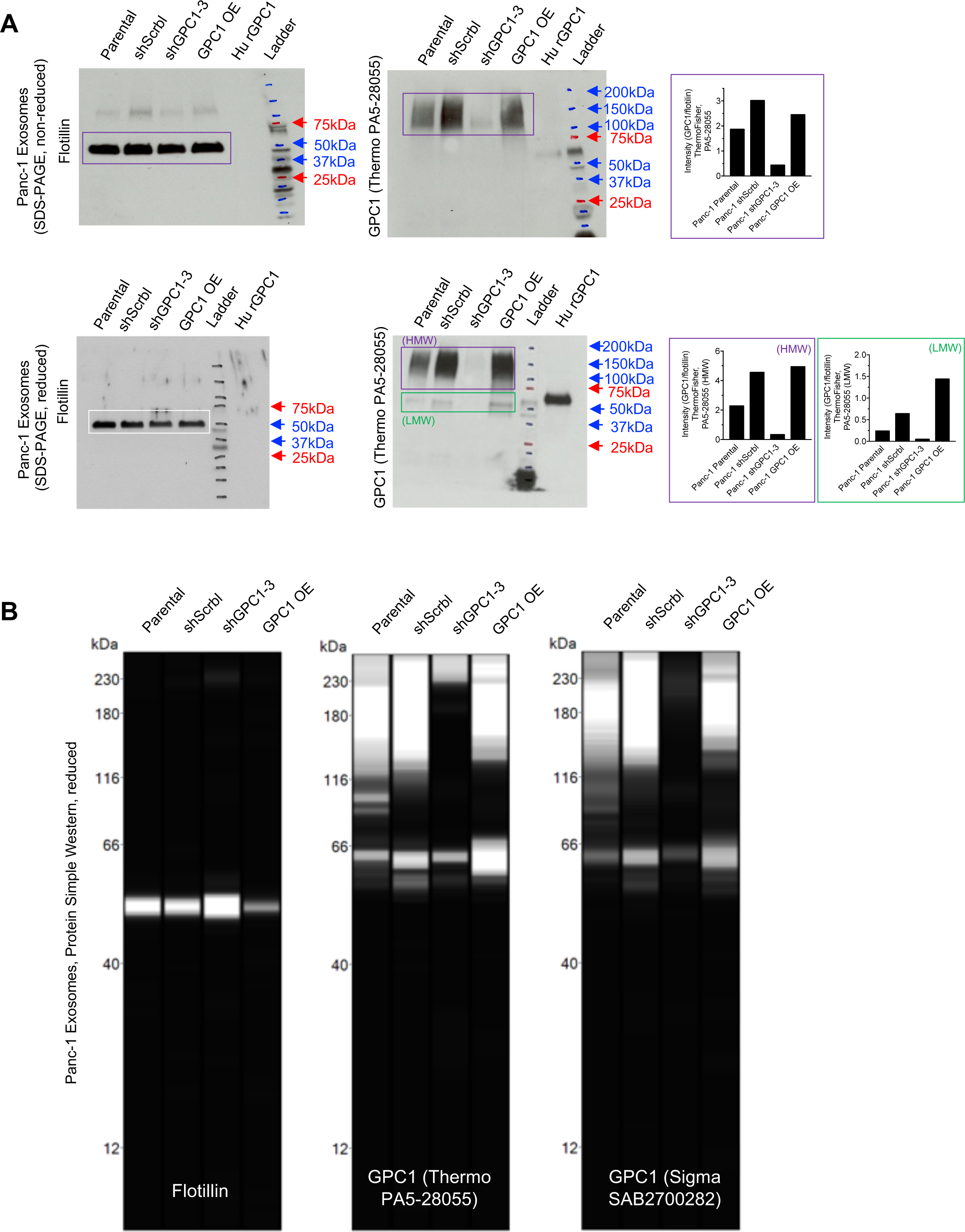
**A**. Western blot analyses of non-reduced and reduced Panc-1 cells-derived exosomes lysates for GPC1 using ThemoFisher PA5-28055. Flotillin is used as loading control. Recombinant human Glypican-1 protein (R&D Systems, 4519-GP-050) was used as positive control (Hu rGPC1). The graphs depict the quantification of GPC1 protein levels (densitometry analysis) relative to flotillin as captured in the indicated boxes. **B.** Protein Simple Western blot analyses of reduced Panc-1 cells-derived exosomes lysates for GPC1 using ThemoFisher PA5-28055 and Sigma SAB2700282 antibodies, flotillin is used as loading control.

**Figure 4.**
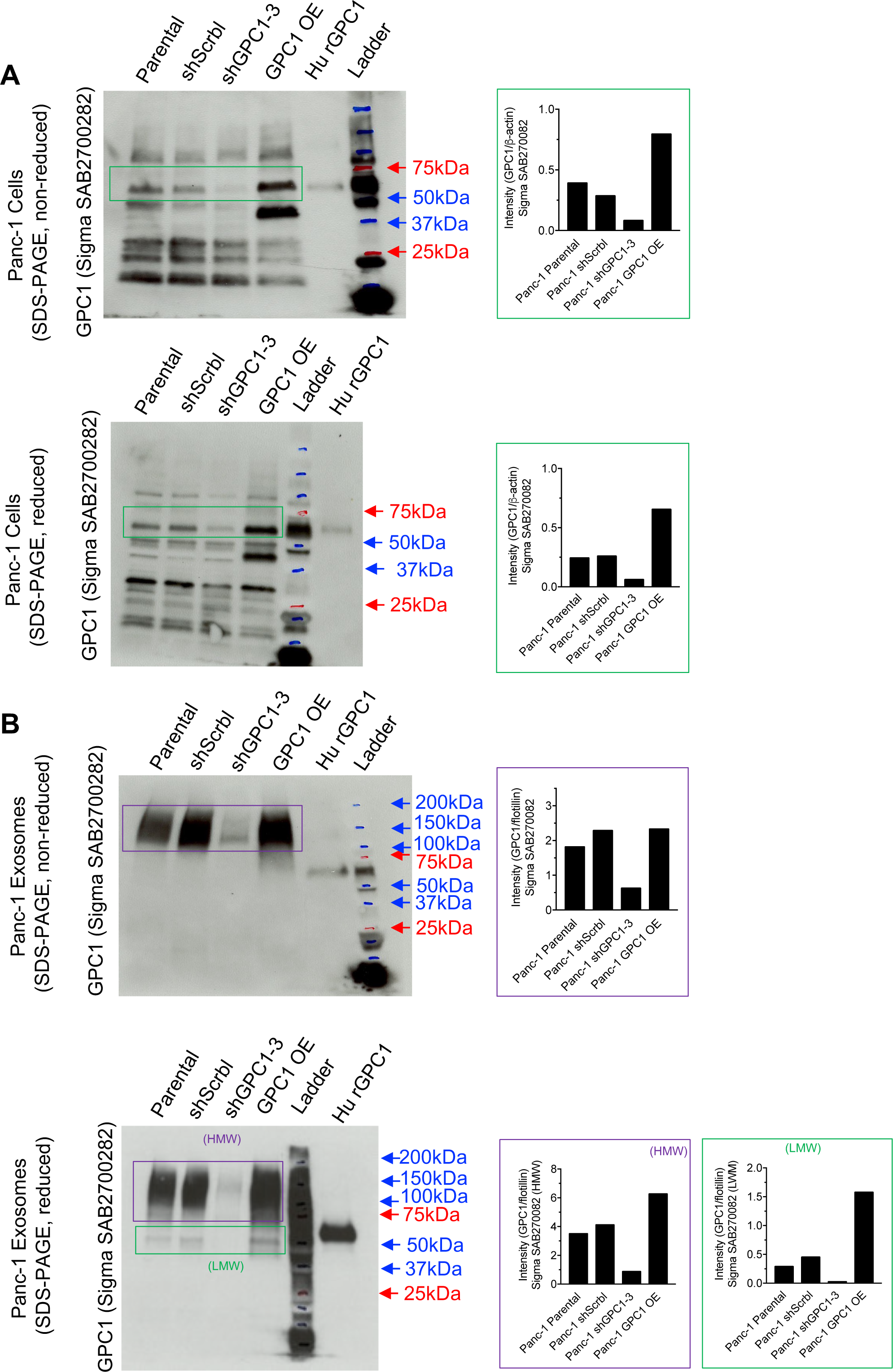
**A.** Western blot analyses of non-reduced and reduced Panc-1 cell lysates for GPC1 using Sigma SAB2700282. β-actin is used as loading control and shown in Figure 2C. Recombinant human Glypican-1 protein (R&D Systems, 4519-GP-050) was used as positive control (Hu rGPC1). The graphs depict the quantification of GPC1 protein levels (densitometry analysis) relative to β-actin as captured in the green boxes. **B**. Western blot analyses of non-reduced and reduced Panc-1 cells-derived exosomes lysates for GPC1 using Sigma SAB2700282. Flotillin is used as loading control and shown in Figure 4A. Recombinant human Glypican-1 protein (R&D Systems, 4519-GP-050) was used as positive control (Hu rGPC1). The graphs depict the quantification of GPC1 protein levels (densitometry analysis) relative to flotillin as captured in the indicated boxes.

Next, we confirmed the specificity of the flow cytometry analyses for GPC1 positive bead-bound exosomes using the Sigma SAB270028 anti-GPC1 antibodies (**Figure 5A-B**), as well as the monoclonal Abnova (MAB8351, clone E9E) antibodies (**Figure 5C**). Additionally, high sensitivity Protein Simple Western blot analyses for Panc-1 cell lysates indicated, using the monoclonal Abnova (MAB8351, clone E9E) antibody, a specific loss of HMW and LMW GPC1 in shGPC1 cells compared to parental and shScrbl controls (**Figure 6A**). These analyses also revealed increased HMW GPC1 in the GPC1 OE cells compared to parental and shScrbl controls and shGPC1 cells (**Figure 6A**). Exosomes analyzed by Protein Simple western blot using the monoclonal Abnova (MAB8351, clone E9E) antibody revealed both HMW and LMW GPC1 in exosomes, with a specific loss of GPC1 in the shGPC1 cells-derived exosomes and an increase in GPC1 in the GPC1 OE cells-derived exosomes compared to parental and shScrbl controls (**Figure 6B**).

**Figure 5.**
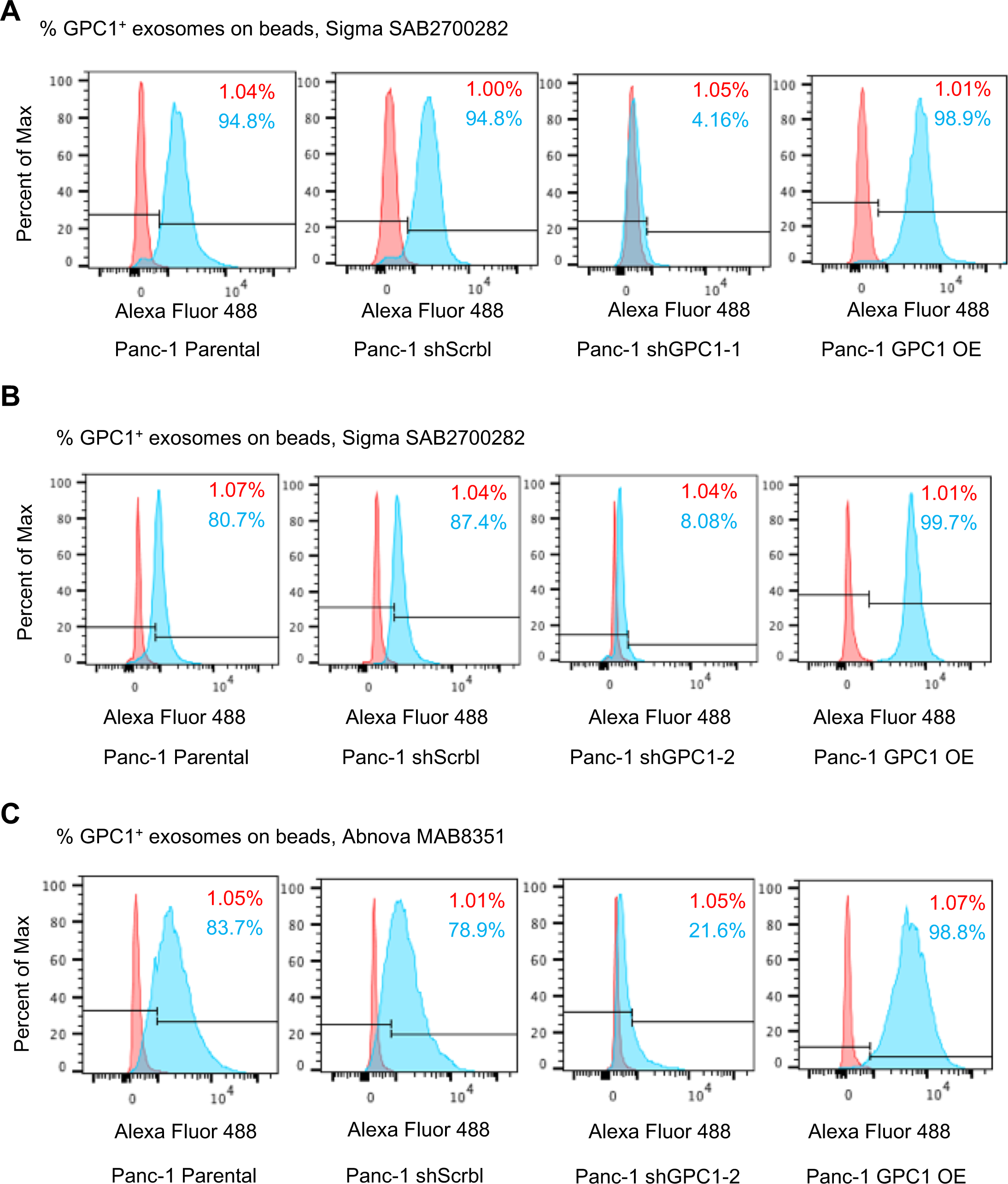
**A-C**. Flow Cytometry of the percentage of GPC1^+^ exosome on beads from Panc-1 cells with anti-GPC1 antibodies from (A-B) Sigma (SAB2700282) or (C) Abnova (MAB8351). Negative control: red peaks, secondary antibody only.

**Figure 6.**
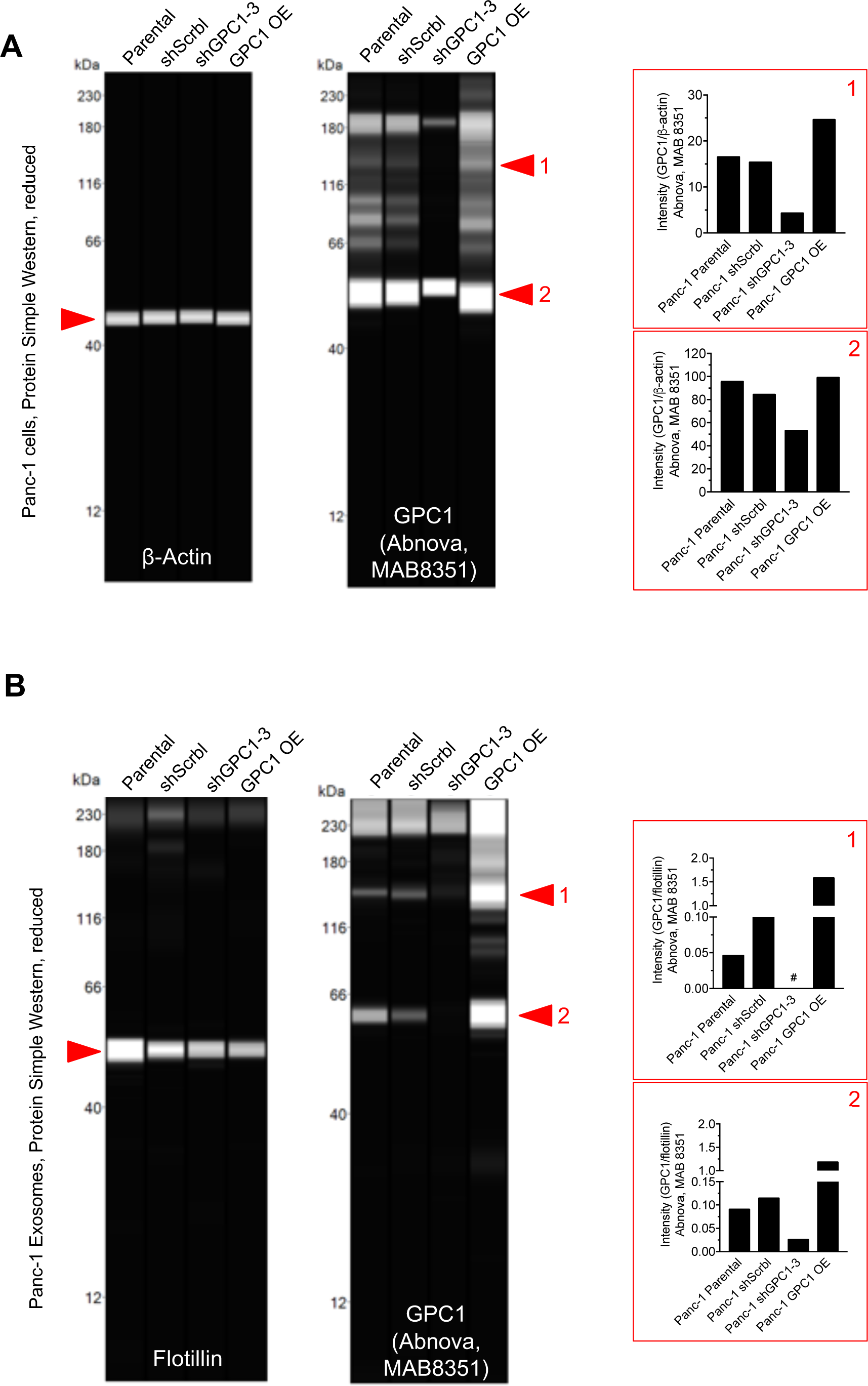
Protein simple western blot analyses of reduced Panc-1 exosomes lysates for GPC1 using AbnovaMAB8351 antibody. Flotillin is used as loading control. The graphs depict the quantification of GPC1 protein levels (densitometry analysis) relative to flotilin. #: no clear peak could be distinguished.

In order to step away from the bead-based analysis of the GPC1 positive exosomes, we analyzed for them using the Flow Nano Analyzer (NanoFCM, www.nanofcm.com), which enables direct visualization of GPC1+ (immunolabeled) exosomes. Using this technique, Sigma SAB2700282 and Abnova MAB8351 anti-GPC1 antibodies revealed a decrease in the percent of GPC1+ exosomes from Panc-1 shGPC1cells when compared to control exosomes (from Panc-1 shScrbl cells), despite a similar number of particle events (exosomes) analyzed (**Figure 7**). In contrast, an increase in the percent GPC1+ exosomes from Panc-1 OE GPC1 cells was observed when compared to control exosomes (**Figure 7**). Gating strategy for these analyses was based on the antibody alone control (see methods).

**Figure 7.**
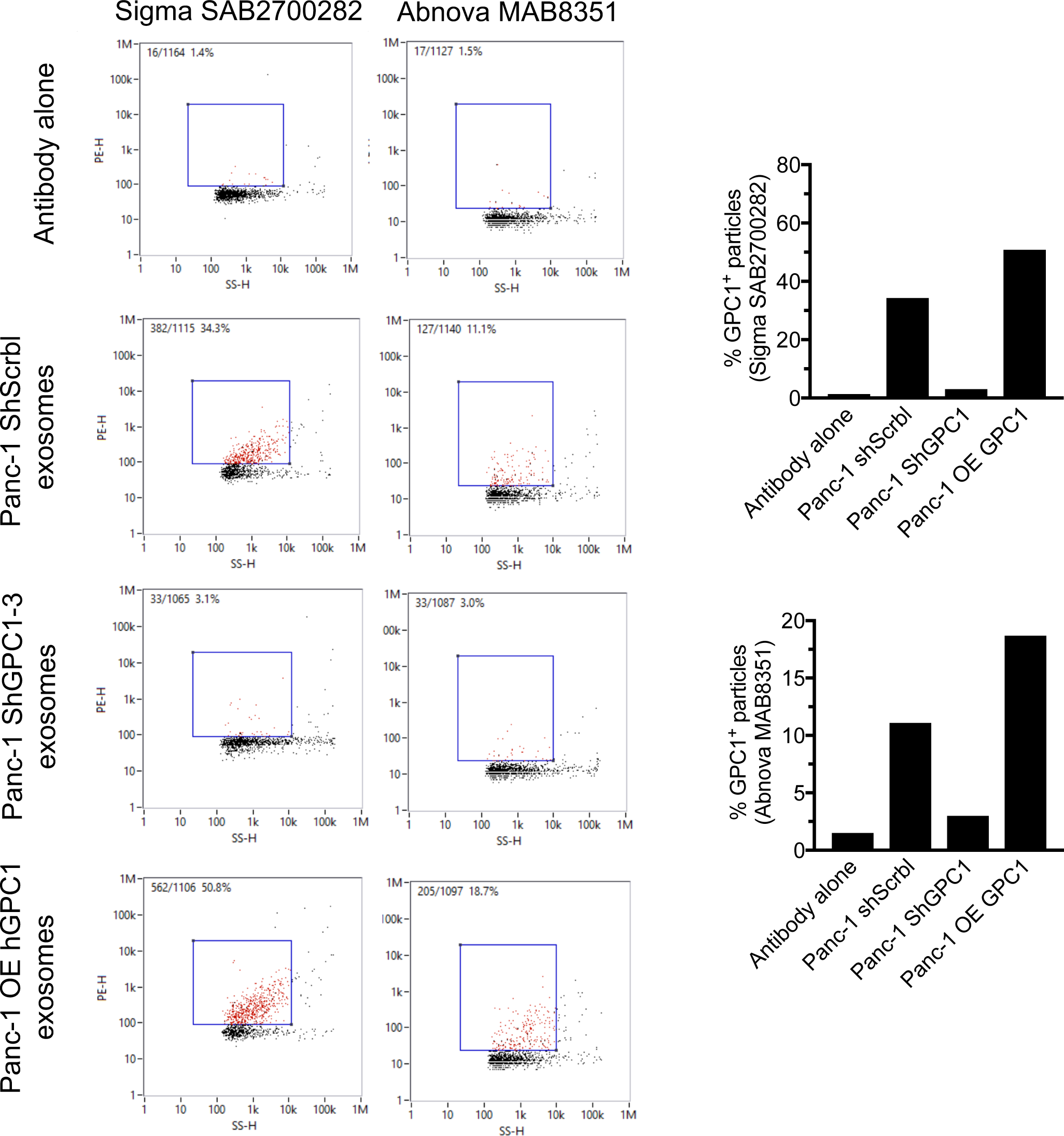
NanoFCM detection of GPC1+ exosomes after the incubation with Alexa Fluro™ 555-conjugated anti-GPC1 primary antibodies (Sigma SAB2700282 or Abnova MAB8351). Antibody only was used as control and to define the gating strategy (blue boxes). The graphs are graphical representation of the percentages listed on each scatter plots.

## Discussion

In the last two years, GPC1 detection on cancer exosomes has been demonstrated by several different independent studies. Such validation included breast cancer cells-derived exosomes and colorectal and esophageal cancer-derived exosomes, in addition to exosomes analyzed from pancreatic cancer cases^2-5,8,14,18-20^ (**Table 1**). Despite employing different methodologies and approaches, GPC1 was reported as elevated in exosomes of patients with different cancer types. Here we provide a detailed methodology, in a stepwise manner, to accompany the findings reported here (**Table 4**). Collectively, 6 different antibodies have been described to identify GPC1 positive cancer exosomes by different laboratories^2-5,8,14,18-20^. We look forward to a productive scientific discourse on the utility of GPC1 positive cancer exosomes in the detection of pancreatic cancer, among others.

## Methods

We provide a detailed protocol in **Table 4** that describes the procedure for exosomes purification and flow cytometry analyses of GPC1 on exosomes. The secondary antibody used was Alexa Fluor 488 goat anti-rabbit (Invitrogen A11008). The aldehyde/sulfate beads were obtained from Invitrogen (A37304). Serum samples were obtained from patients with pancreatic cancer. Serum samples were also obtained from patients with a benign pancreas disease and from healthy donors who had no evidence of acute or chronic or malignant disease, and had no surgery within the past 12 months. The cases were obtained under an IRB-exempt protocol of the MD Anderson Cancer Centre (IRB no. PA14-0154). The flow cytometer used for these analyses was a BD FACS Canto II.

### Cell Culture

Panc-1 cells were obtained from the American Tissue Culture Collection (ATCC) and routinely tested for mycoplasma. 250,000 Panc-1 cells were allowed to adhere to the wells of a six-well plate overnight. Transfection of the shScrbl, 4 shGPC1, and GPC-1 OE plasmid constructs was performed using Lipofectamine 2000, following the manufacturers guidelines. The constructs were purchased from Origene (TF312687) for shGPC1-1 (GGCTGCCTTGCCACCAGGCCGACCTGGA), shGPC 1-2 (CAGCGGTGATGGCTGTCTGGATGACCTCT) and shGPC1-3 (GGAAGGACAGAAGACCTCGGCTGCCAGCT), and GeneCopoeia (EX-Y4957-M56) for GPC1 OE. After two days in selection medium (RPMI supplemented with 10% FBS and 1 Mg/M of puromycin for shScrbl and shGPC1 cells, or 800 Mg/M neomycin for OE GPC1 cells), the cells were single cell-sorted by RFP fluorescence into the wells of a 96-well plate to generate stable cell lines. Single clones with high RFP expression that generated colonies were then expanded, maintained in selection medium, and used for downstream experiments. Notably, we did not detect endogenous RFP fluorescent protein in the exosomes derived from these cells by NanoFCM analyses maybe due to a limited number of molecules included on each exosome.

### Real-time PCR Analysis

One set of Panc-1 cells (Parental, shScrbl, shGPC1 and GPC-1 OE) was chosen and used for each experiment. 200,000 cells of each cell line were plated into a well of a six-well plate. The next day, the cells were harvested for RNA extraction. Three wells were prepared for each cell line at the same time and a well was defined as a biological replicate. Total RNA was extracted using Direct-zol Total RNA miniprep kit (Zymo Research), according to the manufacturer’s directions. Synthesis of cDNA was carried out in a 20µl reaction using High-Capacity cDNA Reverse Transcription kit (Applied Biosystems) according to the manufacturer’s directions. Real-time PCR was performed on an ABI QuantStudio 7 Flex instrument using Power SYBR Green Master Mix (Applied Biosystems). The transcript of interest was normalized to GAPDH transcript levels. Primers for GAPDH: Forward-5’-AATCCCATCACCATCTTCCA −3’; Reverse-5’-TGGACTCCACGACGTACTCA −3’; Primers for GPC1: Forward-5’-TGAAGCTGGTCTACTGTGCTC −3’; Reverse-5’-CCCAGAACTTGTCGGTGATGA −3’. Each reaction included three technical replicates, which were averaged to represent one biological replicate. Statistical analyses (unpaired two-tailed t test) were performed on the dCt of biological replicates and the data were presented as relative fold change (FC) in expression.

### Western Blot analyses

Cell lysates were isolated from Panc-1 cells using cell lysis buffer containing 50mM Tris (PH 7.5), 150mM NaCl, 0.1% SDS, 1% Triton X-100, 1% Sodium Deoxycholate, and supplemented with cOmplete protease inhibitor cocktail tablet (Roche, 11647498001). The lysate was cleared by centrifuging at 4°C, 12,000g for 10 minutes. Total protein concentration was measured using BCA assay and 20µ,g of the non-reduced lysate was mixed with loading buffer. Alternatively, the proteins were reduced by adding b-mercaptoethanol to the loading buffer. The samples were boiled for 10 minutes at 95°C before loading onto the a Mini-Protean TGX Stain-Free 4-15% gel (Bio-Rad, 456-8084) and transferred to PVDF membranes using the Trans-Blot Turbo Transfer System (BioRad, 17001919). Membranes were blocked with 5% milk in TBS-T for 1 hour at room temperature and then incubated with GPC1 primary antibody overnight at 4°C (1:500, Sigma SAB2700282 or 1:1,000 ThemoFisher PA5-280055). The same lysates were run in parallel and the blot was incubated with b-Actin-HRP conjugated antibody for overnight (1:30,000, Sigma A3854). Membranes incubated with anti-GPC1 primary antibody were washed 3 times in TBS-T (10 minutes each) and incubated with anti-rabbit-HRP secondary antibody (1:2,000, Sigma A0545) for 1 hour at room temperature. All blots were then washed 3 times with TBS-T and then developed using Pierce ECL Substrate (Thermo- Scientific, 32106). Exosomes were prepared as previously described^1^ and exosomes lysates were prepared using ProteinSimple RIPA buffer (ProteinSimple, 040-483) supplemented with protease inhibitor (cOmplete protease inhibitor cocktail tablet, Roche, 11647498001, no DTT was added to the sample buffer). To reduced the exosomes lysates, b- mercaptoethanol was added to the loading buffer. Total protein concentration was measured using BCA assay and 20µ,g of the lysate in the 4x Blot LDS sample buffer (ThermoFisher, B0007) were boiled at 95°C for 10 minutes, run on a Mini-Protean TGX Stain-Free 4-15% gel (Bio-Rad, Cat#456-8084), and transferred to PVDF membranes using the Trans-Blot Turbo Transfer System (BioRad, 17001919). Membranes were blocked with 5% milk in TBS-T for 1 hour at room temperature and then incubated with anti-GPC1 primary antibody for overnight at 4°C (1:500, Sigma SAB2700282 or 1:1,000 ThemoFisher PA5-280055) or anti-Flotilin-1 (H-104, 1:100, Santa Cruz, sc-25506). The membranes were washed 3 times in TBS-T (10 minutes each) and incubated with anti-rabbit-HRP secondary antibody (1:2,000, Sigma A0545) or anti-mouse-HRP secondary antibody (1:2,000, Fisher-Scientific, HAF007), for 1 hour at room temperature. All blots were then washed 3 times with TBS-T and then developed using Pierce ECL Substrate (Thermo-Scientific, 32106).

Cell and exosomes lysates were also analyzed by Protein Simple Western blot. To this end, the cell lysates, prepared as described above, was further processed according to the manufacturer’s instructions and DDT was added to the loading buffer. Approximately 200ng of proteins were loaded per capillary, and a 1:500 b-Actin-HRP conjugated antibody (Sigma A3854), or 1:50 dilution of Abnova MAB8351 anti-GPC1 antibody was used. For the exosomes analyses, exosomes (~200ng per capillary equivalent) were directly mixed with the loading buffer (containing DTT) and samples processed as per the manufacturer’s directions. Anti-Flotilin-1 (Invitrogen, PA5-19713) was used at a 1:50 dilution, and the Abnova MAB8351 anti-GPC1 antibody was used at a 1:40 dilution.

### Flow cytometry analyses of exosomes from cells

The flow cytometry analyses of exosomes from cells were developed based on the protocol listed in **Table 4** with minor modifications. The secondary antibody used in this part was Alexa Fluor 488 goat anti-rabbit (Invitrogen A11008). Approximately 5×10^9^ exosomes from each sample were with 10µl of beads and the mixture mixed for 3 hours at 4°C. In the primary antibody binding step, 1-3µ.l of anti-GPC1 antibody (ThemoFisher PA5- 280055, Sigma SAB2700282, or Abnova MAB8351) was used for 1 hour incubation at room temperature, and 3 wash steps were carried out using 200µ1 of 2%BSA/PBS. After the secondary antibody incubation, 3-5 wash steps were carried out using 200µ1 of 2%BSA/PBS. One to 0.5µ1 of secondary antibody can be used for these experiments. A BD LSR Fortessa X-20 Cell analyzer was used to analyze these samples.

### Flow NanoAnalyzer (NanoFCM) Studies

Exosomes were obtained from cells cultured in serum-free RPMI media as previously described^1^ and resuspended in 200µ1 of PBS. Exosomes concentration was measured using the NanoSight LM10 (NanoSight Ltd). Approximately 10^9^ exosomes in 100µ1 of PBS with 3% Triton-T were incubated with 1µl of Alexa Fluor™ 555-conjugated anti-GPC-1 primary antibody (Sigma SAB2700282) or 1µl of a 1:10 dilution (in PBS with 3% Triton- T) of Alexa Fluor™ 555-conjugated anti-GPC-1 primary antibody (Abnova MAB8351) for 10 hours at room temperature while rotating. The Alexa Fluor™ 555 dye was conjugated to anti-GPC-1 primary antibody using a labeling kit (Thermo-Scientific) according to the manufacturer’s directions. The reaction mixture was then purified with Total Exosome Isolation Reagent (Invitrogen, 4478359). Then labeled exosomes were resuspended in 100µL PBS and analyzed using the NanoFCM (NanoFCM Inc. China) as per the manufacturer’s instructions. Gating was defined based on the signal obtained when antibody alone (no exosomes) was subjected to the same procedure as described above.

